# Tactile motor attention induces sensory attenuation for sounds

**DOI:** 10.1101/2021.07.08.451581

**Authors:** Clara Fritz, Mayra Flick, Eckart Zimmermann

## Abstract

Sensory events appear reduced in intensity when we actively produce them. Here, we investigated sensory attenuation in a virtual reality setup that allowed us to manipulate the time of tactile feedback when pressing a virtual button. We asked whether tactile motor attention might shift to the tactile location that makes contact with the button. In experiment one, we found that a tactile impulse was perceived as more intense when button pressing. In a second experiment, participants pushed a button and estimated the intensity of sounds. We found sensory attenuation for sounds only when tactile feedback was provided at the time the movement goal was reached. These data indicate that attentional prioritization for the tactile modality during a goal-directed hand movement might lead to a transient reduction in sensitivity in other modalities, resulting in sensory attenuation for sounds.

## 1 Introduction

A pivotal requirement for successful interaction with the environment is the ability to distinguish sensory stimuli that are produced by ourselves from those that have an external cause. The plethora of examples illustrating this need ranges through the animal kingdom from the electric fish that can dissociate self-and externally produced electric fields (Bell, 2001) to humans who are not disturbed by the retinal motion that is generated by their own eye movements (Wurtz, 2018). Research into these phenomena early on demonstrated that a signal must exist that tells sensory areas about upcoming movements (von Holst & Mittelstaedt, 1950). In consequence, sensory areas alter their receptivity for the sensory effects produced by that movement. The ensuing attenuation of sensory events has been found for tactile (Bays et al., 2006), auditory (Weiss et al., 2011) and visual events (Cardoso-Leite et al., 2010). A comparator model account has been proposed in which the initiation of a movement triggers an efference copy that is used to build up a forward model that predicts the consequences of the movement (Blakemore et al., 2002). If the prediction and the actual sensation match, the strength of that sensation is reduced, resulting in sensory attenuation.

Sensory attenuation has been previously tested for body-related events (Dogge et al., 2019). Predictions focused on action outcomes that are closely related to the direct actions itself. This model has been expanded to account for sensations that are not body-related (Dogge et al., 2019). Thus, explanations and effects of sensory attenuation for external events, like sounds generated by a button-press, must be distinguished from sensory attenuation for self-touch (Bays et al., 2006). Although both phenomena share common elements, evidence suggests that the two effects rely on separate neural mechanisms (Dogge et al., 2019). For instance, sensory attenuation of self-touch is spatially selective for the goal location of the touching movement (Bays et al., 2008; Kilteni et al., 2019; Knoetsch & Zimmermann, 2021). It can be predicted precisely because - unlike in most experiments on attenuation of external events - the position of the tactile sensation exactly matches the position of the touch. Furthermore, the connection between touch and tactile sensation is over-learned across the lifetime whereas the contingency between a movement (e.g. a button press) and an arbitrary external event must be trained to be predictable. However, for external events studies have shown that sensory attenuation occurs even if sounds merely coincide unpredictably with a button press (Bäß et al., 2008; Horváth et al., 2012).

In our study we tested a novel hypothesis concerning sensory attenuation for sounds produced by closed-loop button presses. In closed-loop movements, feedback about the success of a movement is available and informs the actor to stop movement execution.

We assumed that pre-motor attention during these goal-directed hand movements improves tactile sensitivity at the predicted time when the hand will make contact with the desired object. Motor induced attention shifts are known to occur for several movements: For eye - (Deubel & Schneider, 1996) and for pointing (Baldauf et al., 2006) movements, it has been repeatedly demonstrated that visual attention shifts to the goal location of the movement shortly before movement onset. The purpose of that shift is to predict the sensory consequences following movement termination in order to estimate movement success and - in the case of saccade eye movements - to establish visual stability across the movement (Baldauf & Deubel, 2008). A similar predictive attention shift has been found for pointing movements. Deubel and colleagues (1998) demonstrated that for the preparation of a pointing movement to a certain location, the perceptual processing is partially initiated even before movement onset. Similarly, to the attention effects observed at the time of saccades, discrimination performance presented close to the goal object of a pointing movement was higher than when presented at other locations.

Our data suggests that goal-directed movements like button presses induce tactile sensitivity to the moving hand in order to prioritize processing. In consequence, processing in other sensory modalities is decreased. The transfer of attentional resources between the tactile and the auditory modality is especially likely given their functional connectivity. The human somatosensory cortex co-activates with auditory cortex during the processing of vibrations and texture (Butler et al., 2012; Iguchi et al., 2007; Nordmark et al., 2012; Schürmann et al., 2006). An influence of attention on these two cortical systems was already described by Gescheider et al. (1975). When auditory and tactile stimuli were presented individually or simultaneously, the cognitive processing was only impaired for the concurrently occurring stimuli. Thus, the distribution of attention was an important determinant of performance. Also in recent studies, Convento and colleagues (2018) demonstrated that participants were impaired in an auditory frequency discrimination task when they received TMS stimulation over S1 and attended to tactile frequency information.

In this study we raised two major questions: First, is tactile sensitivity more increased at the time the hand makes contact with the button than at the beginning of the movement? It is long known that during arm-movements, tactile gating is responsible for increased tactile thresholds while the movement is ongoing (Chapman et al., 1987). Juravle et al. (2010) tested tactile thresholds while participants had to grasp a computer mouse. They found that tactile sensitivity in the phase after the movement is significantly higher compared to the execution phase of the movement. The threshold values decrease from preparation to execution and increase again from execution to the phase after the movement (Juravle et al., 2010). We tested tactile thresholds during closed-loop button-press movements that are commonly used to measure sensory attenuation of external events (Weiss et al., 2011). Second, does attention processing the tactile sensation during a button press determine attenuation for sounds that are contingent on the button press?

We investigated these questions in a virtual reality setup that allowed us to manipulate the time of the tactile feedback when pressing a button. With our virtual reality setup, we could control the presence of tactile feedback. In studies investigating tactile gating and sensory attenuation an activation of the tactile modality is included, such as participants touching their own hand (Voudouris & Fiehler, 2017) or grasping an object (Juravle et al., 2010). These setups therefore include two tactile sensations, the feedback when touching the object in addition to the probe stimulus delivered by the experimenter. With the help of the virtual reality design, we were able to replace the physical feedback with artificial tactile feedback.

In our study a virtual button was presented in a head-mounted display and tactile feedback was provided via mini-vibrators that were attached to the participants’ index fingers. In Experiment 1, we tested perceived vibration intensity at three different movement phases and in a baseline condition. In Experiment 2, we sought to find out whether the presence of tactile feedback determines sensory attenuation of actively produced sounds. In Experiment 3 we asked whether the putative interaction of tactile stimulus presentation and sensory attenuation for sounds is dependent on active movements.

## 2 General Methods

### 2.1 Participants

A total of 30 study participants took part in Experiment 1 (including one author). For one participant psychometric functions could not be estimated, indicating that the task was not performed correctly. Therefore, the final sample in Experiment 1 consisted of 29 right-handed participants with unrestricted vision or vision correction (age: 18 - 56 years [*M*_age_ = 26.52, *SD* = 9.87], gender: 11 male, 17 female, 1 non-binary).

Moreover, we calculated a Baseline condition for Experiment 1 post-hoc. For this condition we used different participants. Here, 29 right-handed participants with unrestricted vision or vision correction (age: 18 - 52 years [*M*_age_ = 24.46, *SD* = 9.14], gender: 10 male, 19 female) were tested.

In Experiment 2, 25 right-handed participants with unrestricted vision or vision correction took part (age: 19 - 36 years [*M*_age_ = 24.03, *SD* = 3.38], gender: 6 male, 19 female).

Experiment 3 contained 20 right-handed participants with unrestricted vision or vision correction (age: 19 - 64 years [*M*_age_ = 32.55, *SD* = 14.34], gender: 9 male, 11 female).

Participants were recruited in the University Düsseldorf or via social networks. Experiments were approved by the local ethics committee of the Faculty of Mathematics and Natural Sciences of Heinrich Heine University, Düsseldorf (identification number: 757184), and are in accordance with the Declaration of Helsinki. Handedness was assessed using the Edinburgh Handedness Inventory and all participants were classified as right-handers. Participants were compensated with participation hours or remunerated by means of an expense allowance. Informed consent was obtained from all participants.

### 2.2 Power analysis

We conducted a power analysis post-hoc. Fraser and Fiehler (2018) introduced a similar study where participants were asked to do a pointing task with a vibration pad being taped to their index finger. They used a repeated measures ANOVA assessing the effects of time of stimulation (early and late) and the target size.

The interaction in the study of Fraser and Fiehler (2018) showed an effect size of 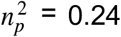. With an alpha level of *α* = 0.05, a Power of 0.8 and *ρ* = 0.55 for a repeated measures ANOVA with three measurements, a total sample size of *N* = 27 for Experiment 1 is required. For a 2×2 repeated measures ANOVA in Experiment 2 a total sample size of *N* = 23 (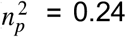, *α* = 0.05, Power = 0.8, *ρ* = 0.55) is required.

A paradigm very similar to ours was used by Gilmeister & Eimer (2007) to study the effects of touch on auditory processing. Since no effect sizes were added by authors, we adapted the number of subjects and also tested 20 subjects in Experiment 3.

As described below, our conclusions depend on many additional tests that are not included in this power analysis. Thus, the identified sample sizes should be considered a minimum for our study.

### 2.3 General materials

The Experiments took place in the same setting with only minimal adjustments. Participants in the experiments were asked to press a button in a virtual reality environment, which was presented to the participants through VR goggles (Oculus Rift Development Kit 2).

The VR goggles included a horizontal and vertical field of view of 100° and a refresh rate of 60 Hz. In the head mounted display, participants saw a blue/green virtual button in front of a dark background. Hand movement of participants was captured with a motion sensor (Leap Motion, Orion V 2.3.1+31549 sampling at 60 Hz with a field of view of 140 x 120°). A virtual hand model was shown that moved synchronously with the real hand. Vibrotactile stimuli were presented via mini-vibrators that were attached to the right index fingers of the participants. The vibrotactile stimuli were controlled by an Arduino Nano microcontroller ATmega328 operating at 5 Volt. The experiments were run on a MacBook Pro (Retina, 15-inch, 2015).

## 3 Experiment 1

### 3.1 Procedure in Experiment 1

In Experiment 1, participants had to perform a goal-directed hand movement to press the virtual button with their right hand. In each trial they made an arm movement starting with the right hand and forearm held up and moved downwards to press the button (see Figure 1). The virtual button went down 5° when pressed.

**Figure 1.**
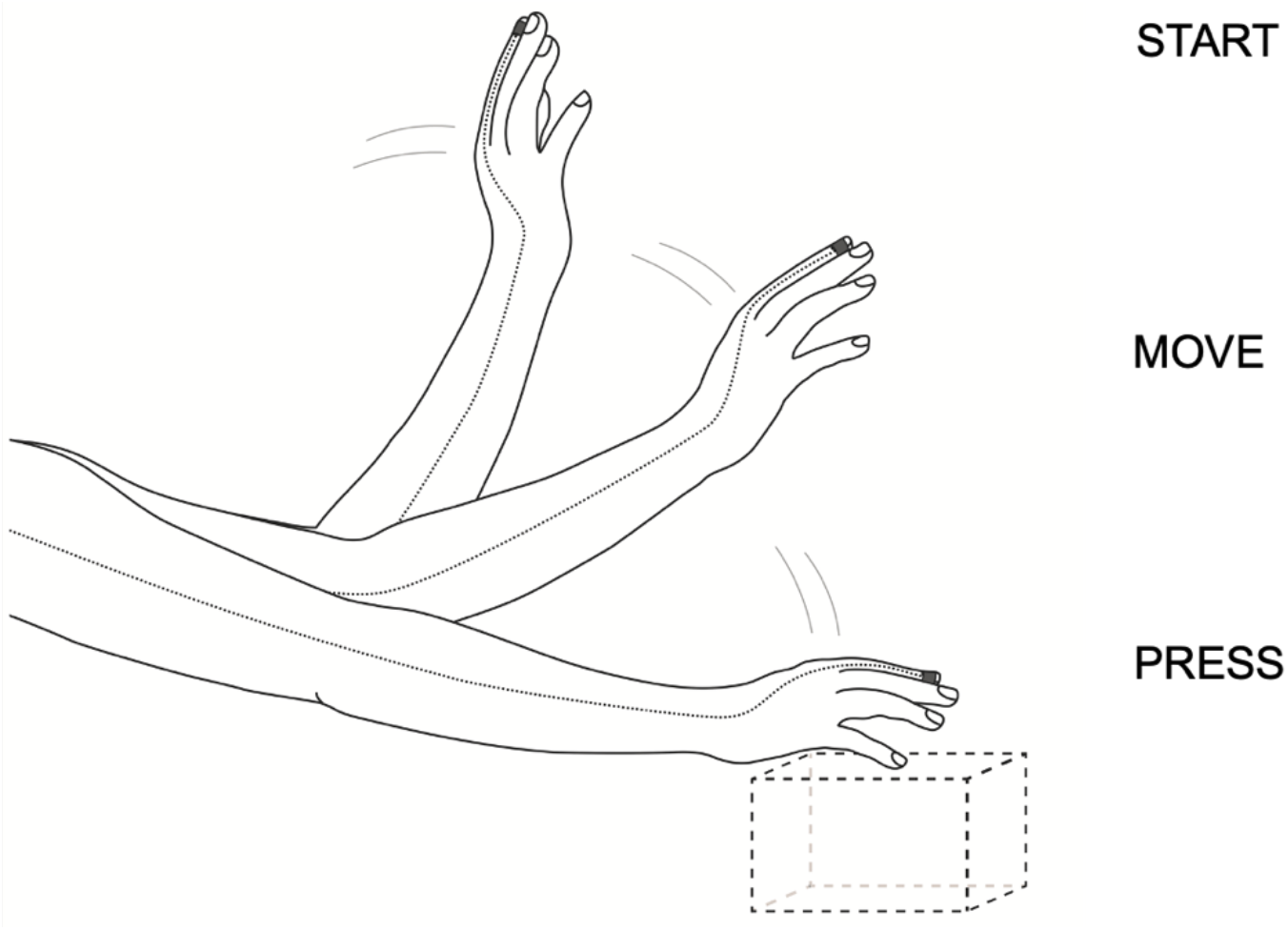
Experimental setup. Participants were asked to perform a goal-directed hand movement to press a virtual button. During the movement the hand starts perpendicular to the button (Start), is then moved down towards the button (Move) and ends the movement with a button press (Press). The mini-vibrators attached to the right index finger are shown as black dots and their cables as grey lines. Participants experienced a tactile stimulation on the right moving index finger either during Start, Move or Press hand movement time (Experiment 1). A comparison stimulation was delivered 700ms after the button press to the resting index finger of the left hand. The virtual button is presented with a dotted rectangular box.

Each trial started with the presentation of a “*Ready*” message that was displayed on the left side of the screen center. After 500 ms, it was replaced by a “*Set*” message which was presented for 500 ms. Participant’s task was to press the button at the corresponding “Go” time, i.e. 500 ms after the appearance of the “Set” message. A vibrotactile stimulus was presented to the right index finger of the moving hand either before the start of the movement (Start), during the movement (Move) or when the button was pressed (Press). The first movement phase included all trials with tactile stimulations that occurred before the go-signal, the second movement phase included all trials in which tactile stimulation occurred equal to or after the go-signal and the third phase included all trials in which tactile stimulation occurred at the exact same time the button was pressed in virtual reality. These presentation times of the vibrotactile stimulus were randomized across trials, so the vibration could either occur during start, move or press phase. The vibration on the index finger of the right moving hand was constant in each trial (50% of the maximum vibration intensity of 5 Volt). After the button was pressed, a comparison vibration was delivered 700 ms later to the index finger of the resting left hand. The comparison vibration intensity varied randomly across trials between 20-40% and 60-80% of the maximum value of 5 Volt (in 6 equidistant steps). For a vibration intensity of 20%, 30% and 40% the vibromotor was driven with a voltage of 1,7 V, 2V and 2,3 for 300 ms each. For a vibration intensity of 60%, 70% and 80% the motor was driven with 2.9 V, 3.2 V and 3.5 V. As the vibration was strongest with a control of 4.1 V and no longer perceptible under 1.1V, we chose these values as the start and end points of vibration control. The movement phase as well as the vibration strength was chosen randomly. After each trial, participants had to decide which vibration was perceived stronger by using a foot pedal (UPC ECS-PPD-F) placed under the table. Pedal pressing was counterbalanced between participants with either pressing the right pedal when the first vibration felt stronger and the left button when the second vibration felt stronger or vice versa. Then, after giving an answer, the next trial started immediately. A total of 180 trials were conducted in the first session (60 trials for each of the three movement phases; ten trials for each vibration intensity of the comparison vibration per time point).

In the Baseline Condition of Experiment 1 participants were required to keep their arm stationary. They were seated in front of a table with the VR goggles on and were asked to distinguish between the same stimuli levels presented in the movement conditions. Stimuli were presented with a temporal interval duration of 750 ms. The foot pedal was used to decide which vibration felt stronger.

### 3.2 Data analysis

For Experiment 1, we analyzed offline when the vibration occurred with regard to hand movement position. In total we differentiated between three hand movement phases.

The first movement phase included all trials with tactile stimulations that occurred before the go-signal, the second phase included all trials in which tactile stimulation occurred equal to or after the go-signal and the third phase included all trials in which tactile stimulation occurred concurrently with the button press. On average, the number of trials per movement phase were distributed as followed: Start *M* = 71.18, *SD* = 7.2; Move *M* = 51.38, *SD* = 4.74 and Press *M* = 65.54, *SD* = 4.02. For all experiments, data were averaged for tactile/auditive intensities within each participant for each of the movement phases and Baseline. Afterwards all data were then fitted by a cumulative gaussian function. The point of subjective equality (PSE) represents the magnitude at which the probe vibration (tactile stimulation) is perceived as stronger than the comparison vibration/tone on fifty percent of the trials.

To identify whether there were significant differences between the PSEs and JNDs during the three movement phases in Experiment 1, a one-way repeated measures ANOVA was used. The data that supports the finding of the study are available at: https://osf.io/edt74/.

### 3.3 Results Experiment 1

In Experiment 1 a tactile stimulation on the right index finger was felt during one of the three hand movement phases: Start, Move and Press. Data for the three movement phases were fitted by cumulative gaussian functions individually for each participant. Psychometric functions of one exemplary participant are shown in Fig. 2. A one-way repeated measures ANOVA was conducted to identify whether the PSEs for the three hand movement phases differ significantly from each other (*M*_Start_ = 5.16, *SD* = 10.64, *M*_Move_ = 8.73, *SD* = 8.73; *M*_Press_= 11.96, *SD* = 9.09). There was a statistically significant difference between all three movement phases (*F*(2, 56) = 8.23, *p =* .002, 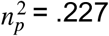). A post hoc analysis (tested against a Bonferroni-adjusted alpha level of 0.05/3) for the repeated measures ANOVA of PSEs revealed that there was a significant difference between the Start and Press movement phases (*MD* = 6.80, *SEM* = 2.03, 95% CI [2.63, 10.96], *p* < .001). No significance was found between Start and Move (*MD* = 3.57, *SEM* = 1.56, 95% CI [0.92, 7.37], *p* = .113) and between Move and Press (*MD* = 3.27, *SEM* = 1.37, 95% CI [−0.56, 7.715], *p* = .179).

**Figure 2.**
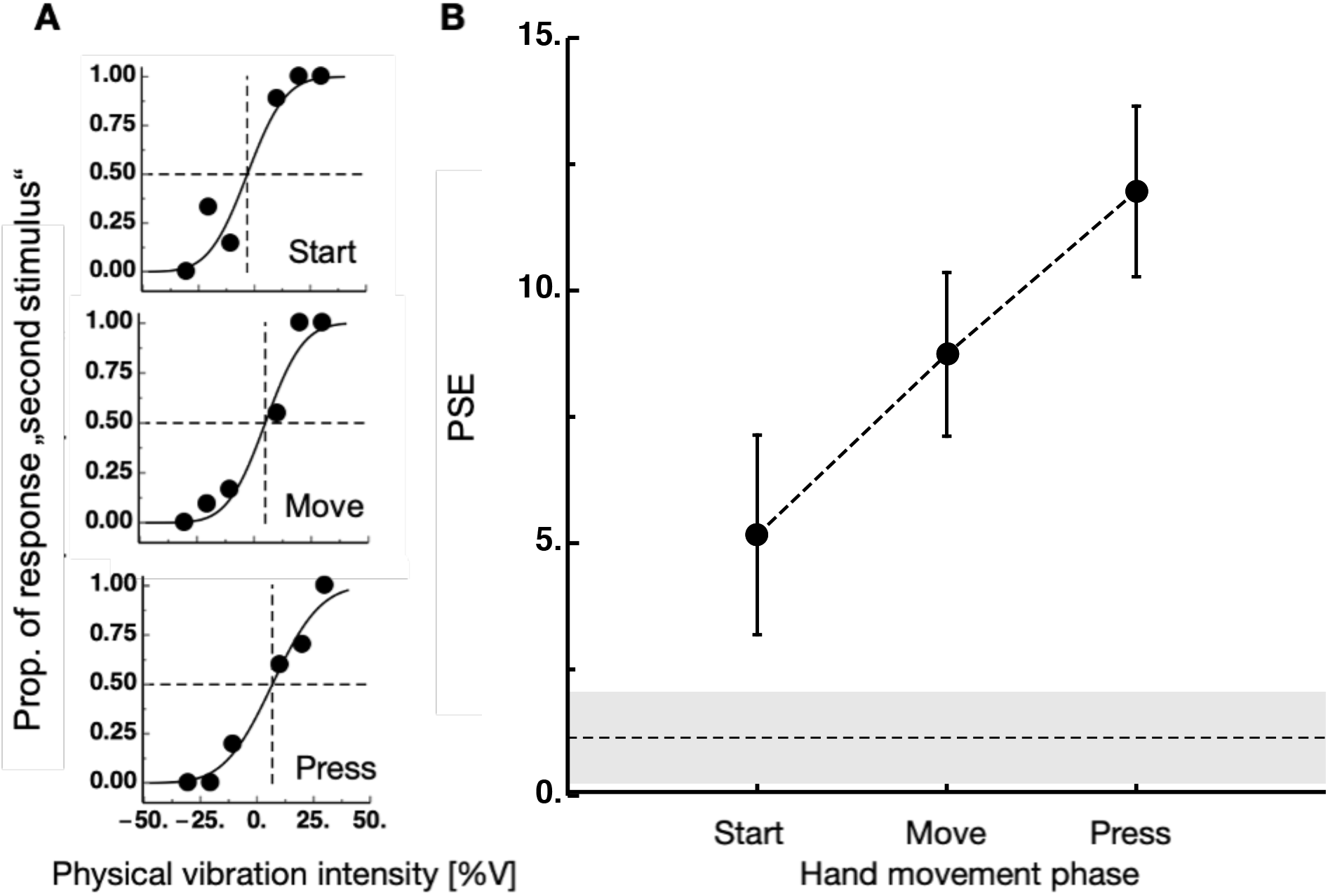
Results of Experiment 1. (A) Psychometric functions of an example participant for the three movement phases. Proportions of second stimulus (corresponding to intensity at resting hand was stronger) are shown against physical vibration intensity. 50% of physical vibration intensity refers to a value of 0 here. (B) Average perceived vibration intensities for the three hand movement phases. The abscissa shows the deviation from the standard stimulus (positive numbers represent overestimation of vibration intensity). Error bars represent S.E.M. Black dotted line with greyish background shows the results for the Baseline condition.

Moreover, we compared all three conditions to the Baseline (*M*_Baseline_ = 51.15, *SD* = 4.92) with Bonferroni corrected paired t-tests (*alpha* = 0.05/3). We found no significant effects between Baseline and the Start phase (*t*(56) = 1.843, *p* = 0.071). We found significant effects between Baseline and Move (*t*(56) = 4.077, *p* < .001, *d* = 1.071) as well as Baseline and Press (*t*(56) = 5.632, *p* < .001, *d* = 1.479).

We also analyzed JNDs with a one-way repeated measures ANOVA (*M*_Start_ = 18.27, *SD* = 17.88; *M*_Move_ = 12.89, *SD* = 16.45; *M*_Press_ = 16.23, *SD* = 17.36). No significant main effect for this ANOVA was found here (*F*(2, 56) = 1.757, *p* = .182, 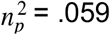).

## 4 Experiment 2

### 4.1 Procedure in Experiment 2

Experiment 2 was divided into two conditions, called ‘no-tactile’ and ‘tactile’. It was randomized whether participants first started with the ‘no-tactile’ or ‘tactile’ condition. Both experimental conditions were conducted in the same virtual reality setup as Experiment 1. Again, participants were asked to perform a goal-directed movement to press the virtual button with the right hand within VR.

In the ‘no-tactile’ condition, a reference tone (MacBook sound ‘Funk’) was presented either during the start of the movement or when pressing the button. As the generation of a tone when button pressing seems quite natural, we assumed that participants perceived the tone as self-generated when pressing the virtual button. We only initiated a second movement time here (during start of the movement). We reasoned that for start as well as move phase no effects of sensory attenuation are to be found.

The tone was presented with 62.3 dB (50% of maximum intensity of the MacBook) through headphones. 700 ms after finishing the goal-directed movement, a probe tone was played, either with 80 - 60% or 40 - 20% of the maximal auditive intensity (67.2, 65.1, 63.6, 58.6, 55.1 or 58.6 dB). We based the individual sound volumes on the MacBook’s loudness gradations. We then evaluated the associated decibel values using a decibel meter. Participants were asked to estimate which tone (reference vs. probe) was louder by pressing either the right or left button of the foot pedal placed on the floor. It was randomized between participants whether the first or second tone was assigned to the right side of the foot pedal. The timing of the reference tone (Start vs. Press movement time) as well as the volume of the probe tone (80 - 60% or 40 - 20%) was randomized. A total of 120 trials was conducted in the condition ‘no-tactile’ (60 trials for each of the two movement phases; ten trials for each auditory intensity per time point).

In the condition ‘tactile’ of Experiment 2 the main part of the test procedure remained similar to the condition ‘no-tactile’. Each trial started 500 ms after a *“Set”* message and a probe tone was played either before or after the goal-directed movement. However, a mini-vibrator was attached to the participant’s right index finger. Each time the button was pressed within VR, participants felt a vibration on the right index finger of their moving hand. More precisely, in 100% of trials in condition ‘tactile’ a stimulus was delivered to the right moving index finger during the button press. In 50% of trials a tone was played when starting the movement and in 50% of trials the probe tone was played when pressing the button. Headphones prevented auditory perception of the vibration sound.

A total of 120 trials was conducted in the condition ‘tactile’ (60 trials for each of the two movement phases; ten trials each for each auditory intensity per time point). Presentation time of the stimulus and auditory intensity were randomized across trials.

For experiment 2 we used a 2 x 2 factorial design to perform an ANOVA for repeated measures to find significant differences between the PSEs and JNDs of ‘no-tactile’ and ‘tactile’.

### 4.2 Results Experiment 2

In half of the trials in Experiment 2 a tone was played when subjects pressed the button and in the other half the tone was played when they started the movement. Additionally, in the condition ‘tactile’, a tactile stimulation was felt on the moving index finger when subjects pressed the button. In the other condition ‘no-tactile’, a tactile stimulation was not felt. A 2 x 2 repeated measures ANOVA with tactile stimulation (‘tactile’ vs. ‘no-tactile’) and hand movement time (Start vs. Press) as factors and the PSEs of the perceived auditory intensities as the dependent variable was conducted (‘tactile’: *M*_Start_ = −0.81, *SD* = 9.04; *M*_Press_ = −7.58, *SD* = 12.37; ‘no-tactile’: *M*_Start_ = −1.81, *SD* = 12.31; *M*_Press_ = −2.84, *SD* = 10.32). The two main effects for tactile stimulation (*F*(1, 24) = 1.302, *p* = .265, 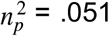) as well as hand movement time (*F*(1, 24) = 3.839, *p* = .062, 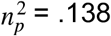) showed no significance (see Figure 3). The interaction between the hand movement time * stimulation showed a significant difference (*F*(1, 24 = 4.71, *p* = .04, 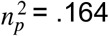). A post hoc analysis (tested against a Bonferroni-adjusted alpha level of 0.05/2) revealed that this significance is to be found between the condition of Start and Press in the tactile condition (*MD* = 6.77, *SEM* = 2.39, 95% CI [0.148, 13.39], *p* = .021). No significance was found between the other conditions (*p* > .176).

**Figure 3.**
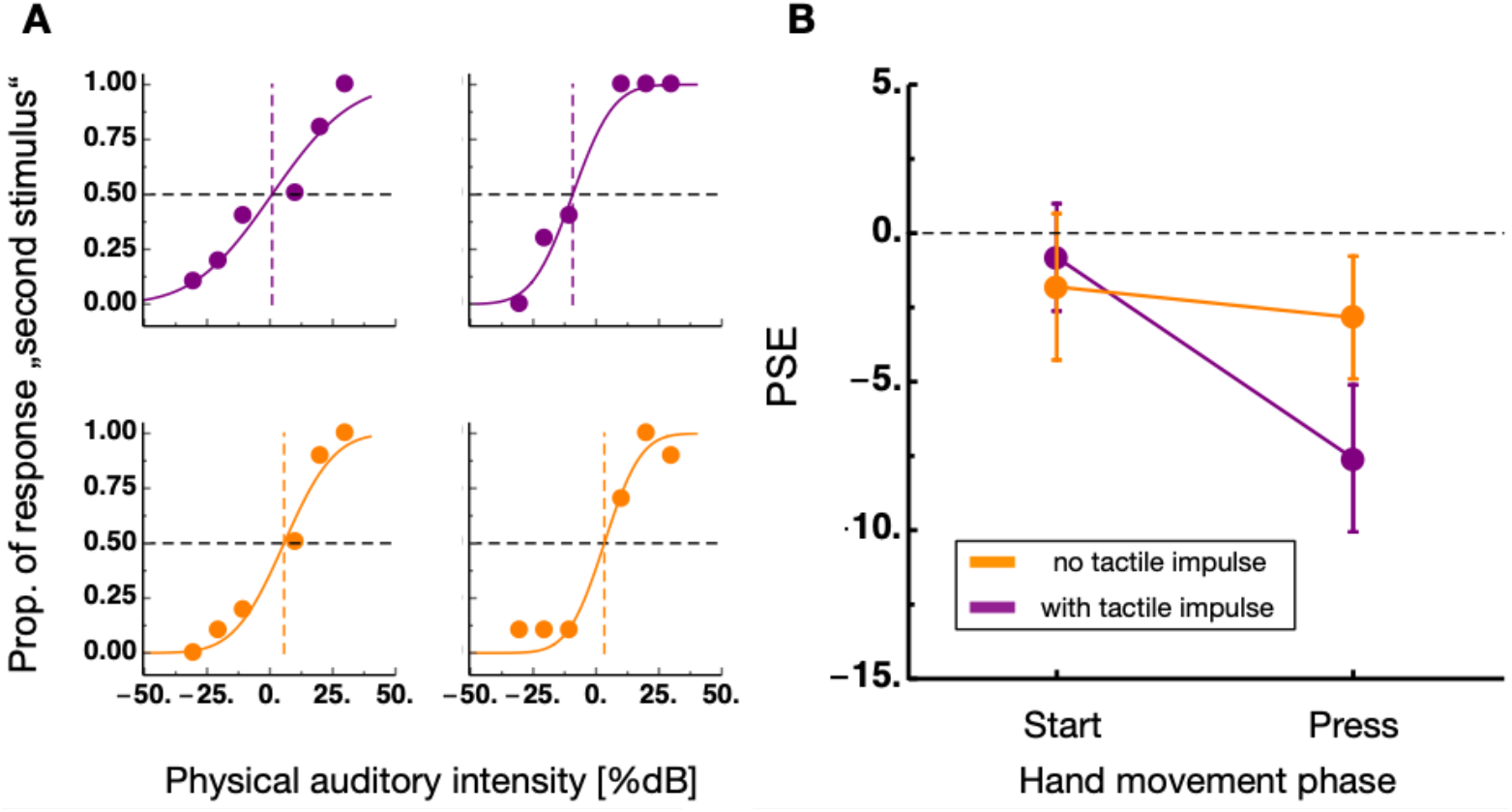
Results of Experiment 2. (A) Psychometric functions of an example participant for the two conditions. Proportions of response for second stimulus are shown against physical auditory intensity. (B) Average perceived sound intensities from session with (shown in purple) and without (shown in orange) tactile stimulation for sounds presented at the Start and the Press time of the movement. The abscissa shows the deviation from the standard stimulus (positive numbers represent overestimation of auditory intensity). Error bars represent S.E.M.

We also analyzed JNDs with a repeated measures 2 x 2 ANOVA with tactile stimulation (‘tactile’ vs. ‘no-tactile’) and hand movement time (start vs. press) as factors (‘tactile’: *M*_Start_ = 28.02, *SD* = 30.9; *M*_Press_ = 22.35, *SD* = 20.18; ‘no-tactile’: *M*_Start_ = 19.08, *SD* = 12.39; *M*_Press_ = 15.71, *SD* = 13.51, 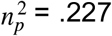). The main effect of hand movement time (*F*(1, 24) = 1.519, *p* = .230, 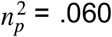) was not significant. However, there was a main effect for the tactile stimulation (*F*(1, 24) = 6.24, *p* = .020, 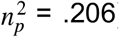). No significant interaction was found (*F*(1, 24 = 192, *p* = .665, 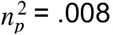).

## 5 Experiment 3

### 5.1 Procedure in Experiment 3

To identify whether the putative interaction of tactile stimulus presentation and sensory attenuation for sounds is dependent on the active movement, participants did not perform the button pressing movement during Experiment 3. Participants were placed in front of a table with headphones on and were told to keep their hands rested on the table without any movement. During the test phase they heard two different tones via headphones. The first tone was played with 50% of the maximum auditive intensity of the MacBook Pro. The second tone, as a comparison tone, was presented 700ms later with either 80%, 70%, 60% or 40%, 30%, 20% of the maximum auditive intensity. After each trial participants had to decide which tone was perceived louder by using a foot pedal placed under the table. Participants had to press the right button of the foot pedal, when the first tone was perceived as louder or the left button when the second tone was perceived as louder (or vice versa). After entering the answer, the next trial started immediately. In the first condition, no tactile stimulation was presented during the discrimination task (‘no-tactile’). In the second condition (‘tactile’), a mini-vibrator was attached to the participants right index finger. A vibration with 70% of maximum intensity was felt on the index finger paired with the first tone. The order of ‘tactile’ vs. ‘no-tactile’ was randomized between participants.

### 5.2 Results in Experiment 3

To analyze data in Experiment 3 a t-test for repeated measures was used. Data was divided into ‘tactile’ (a tactile stimulation was felt during the sound discrimination) and ‘no-tactile’ (absence of tactile stimulation during sound discrimination). A t-test for repeated measures revealed no significant differences in the PSEs for ‘tactile’ (*M* = 53.895, *SD* = 7.555) and ‘no-tactile’ (*M* = 51.15, *SD* = 6.93): (*t*(19) = 1.3, 95% CI [−1.67, 7.16], *p* = .208). Also a repeated measures t-test for JNDs showed no significant effects for the difference between ‘tactile’ (*M* = 14.03, *SD* = 8.84) and ‘no-tactile’ (*M* = 14.55, *SD* = 9.38): *t*(19) = −0.188, 95% CI [−6.31, 5.27], *p* = .853). The results for Experiment 3 are presented in Figure 4.

**Figure 4.**
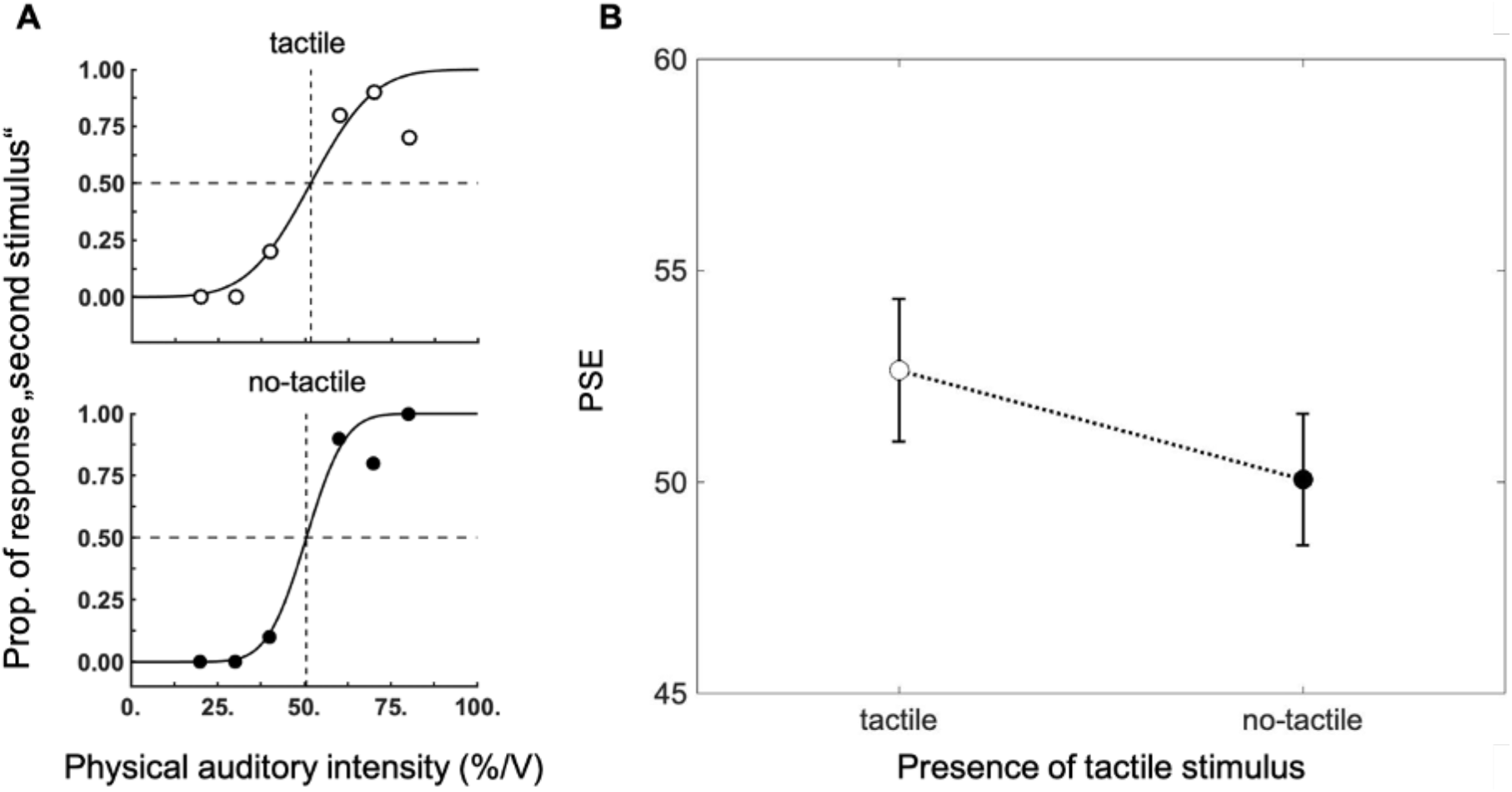
Results of Experiment 3. (A) Psychometric functions of an example participant for the two conditions. Proportions of response to the second stimulus are shown against physical auditory intensity. (B) Average perceived sound intensities from session with and without tactile stimulation. Error bars represent S.E.M.

## 6 Discussion

In this study we investigated the underlying processes that could be responsible for sensory attenuation of self-generated sounds during closed-loop movements. In Experiment 1 we asked if in goal-directed button pressing movements, tactile sensitivity is increased at the time the hand makes contact with the button. We found that apparent tactile stimulus intensity was higher at the time when the hand reached the movement goal than before or during the movement. This finding indicates that tactile attention increases at the time when the hand reaches the goal object and transiently improves tactile sensitivity. Moreover, we introduced a no-movement Baseline condition here which showed significantly lower PSE values than the Move and Press condition. This rules out that effects could be driven by the presentation time of stimuli, as the first stimulus appeared earlier in relation to the second stimulus in the Start and Move phases compared to the button press.

Previous studies found a reduction of tactile sensitivity during arm movements, a phenomenon known as gating (Chapman et al., 1987). The phenomenon has been described frequently for reaching and grasping movements (Colino & Binstead, 2016; Colino et al., 2017). Tactile gating demonstrates that tactile stimuli on the hand are harder to detect during movement (Buckingham et al., 2010; Colino et al., 2017). Usually, this phenomenon starts shortly before movement initiation and builds up again towards the movement goal (Juravle et al., 2010) and tactile sensitivity in the phase after the movement is significantly higher compared to the execution phase of the movement (Voudouris & Fiehler, 2021). Suppression for probed body parts that did not make contact with the goal object is significantly higher (Colino et al., 2014). Suppression of tactile information is clearly driven by task-relevance. As Manzone et al. (2018) showed, suppression during movement varied with task relevance so that targeted movements showed less suppression (Manzone et al., 2018, Colino et al. (2014). In our study, we provide tactile feedback directly to the pressing finger making the tactile stimulation essential to end the closed-loop the button press. Thus, task-relevancy of the tactile feedback in our study is maximized, explaining the absence of suppression. As we conducted our study in a virtual reality setup, no other tactile sensation could confound effects. In other studies, such as Juravle et al. (2010) or Voudouris and Fiehler (2017) this confounding was less controlled. In the study of Juravle et al. (2010), participants moved their hand from a computer mouse (start position) to a goal mouse (goal position) at the end of the table. They found that tactile thresholds increased in the moment the hand picked up the object. Though, tactile thresholds were measured while the finger made physical contact with the object. The sensation of pressure might have reduced the apparent intensity of the tactile impulse.

In Experiment 2 we wondered whether the boost of tactile sensitivity we observed in Experiment 1 would lead to sensory attenuation for sounds. Participants judged the intensity of the sounds in two conditions: Virtual button presses were either accompanied by a tactile stimulation or not. As stated above, for saccades, Deubel and Schneider showed (1996) that at the time of saccade onset, attention is bound to the target position. However, a recent study showed that attentional shifts preceding saccades that are executed to an extinguished target allow attention to spread (Szinte et al., 2019). Thus, in the absence of the visual target, attention is no longer bound to the goal location of the saccade. Following this analogy, we assumed that only in the presence of tactile stimulation, attention would be bound to the tactile modality. Indeed, we found sensory attenuation for sounds only when virtual button presses produced a tactile stimulation.

In principle, this interaction could also occur without the movement. However, sensory attenuation of sounds has been shown to occur only for active movements (Timm et al., 2014). In Experiment 3 we tested auditory perception in the presence and absence of tactile stimuli without movements. Here we found no significant differences, supporting the effects of Timm and colleagues (2014) that the hand movement was a prerequisite for the given effects of attentional shift.

The occurrence of sensory attenuation in the auditory domain is consistent with current literature findings (Mifsud & Whitford, 2017; Timm et al., 2014). However, we believe that the boost of tactile sensitivity observed in Experiment 1 could be an indicator for sensory attenuation in Experiment 2. We theorize that the reaching of the movement goal, e.g., touching the button, is prioritized in the movement of button press. Therefore, attention is bound to the touching finger at this exact moment. In line with our theory, in the condition in which no tactile stimulus was presented and no attentional resources were required for its processing, sensory attenuation was virtually absent. Attention shifts are known to amplify neural responses and perceived intensity (Desimone & Duncan, 1995; Posner & Dehaene, 1994; Posner & Petersen, 1990), thus explaining the increase in tactile and the decrease in auditory sensitivity. Why would attentional prioritization of the tactile modality create an imbalance between the tactile and the auditory modalities? This linkage might be explained by the close neural connectivity of both modalities. Coactivation of the somatosensory and the auditory cortex have been reported previously (Butler et al., 2012; Convento et al., 2018; Iguchi et al., 2007; Nordmark et al., 2012; Schürmann et al., 2006) and an influence of attention on these two cortical systems was already described by Gescheider and colleauges (1975).

On the one hand, a putative limitation of our study might consist in the combination of the virtual button and the presentation of the tactile impulse, which subjects did not experience before participating in the experiment. On the other hand, with this setup we could successfully replicate the sensory attenuation effect. The idea that a sensorimotor contingency between the button press and the ensuing sensory effect must be established before the start of the experiment is based on a theoretical assumption about sensory attenuation that might be incorrect. Three EEG studies found that prediction of sensory consequences is not necessary for sensory attenuation of sounds to occur. When self- and externally generated stimuli were presented within the same experimental block, Baess et al (2011) found that N1 suppression for self-generated stimuli was even stronger. Furthermore, a study also showed that sensory attenuation occurs when stimuli are not predictable but merely coincide with a button press (Horváth et al., 2012; Lange, 2011). In Experiment 2, we showed that in the absence of tactile stimulation, no attenuation was observed. In itself the visual observation of the hand pressing the button has no influence on the reduced sensory intensity of sounds. A visual-tactile interaction might influence the observed effect. The most plausible reason why this should occur is that the visual hand motion allows to predict the time when the tactile stimulus will be applied.

### Conclusion

In conclusion, we found sensory attenuation for sounds when tactile feedback was provided during a button-press movement but not when there was an absence of tactile stimulation. Our data suggest that during goal-directed hand movement’s attention transiently boosts tactile sensitivity at the time the hand reaches the goal object. This increase might be responsible for an imbalance between the tactile and the auditory domain, leading to reduced attentional resources in the latter and thereby to sensory attenuation for sounds.

## Acknowledgements

This research was supported by the European Research Council (project moreSense, grant agreement 757184) and by the Deutsche Forschungsgemeinschaft (DFG, ZI 1456).

## Author Contributions statement

EZ developed the study concept. All authors contributed to the study design. Testing and data collection were performed by CF and MF. CF performed the data analysis and interpretation under the supervision of EZ. All authors contributed to writing the several drafts of the paper and approved the final version of the manuscript for submission.

## Data statement

The data of the study are available at https://osf.io/edt74/.

## Author Note

An earlier version of this manuscript was published as a bioRxiv-preprint, available at https://www.biorxiv.org/content/10.1101/2021.07.08.451581v1

